# Omicron variant escapes therapeutic mAbs contrary to eight prior main VOC

**DOI:** 10.1101/2022.01.03.474769

**Authors:** Céline Boschi, Philippe Colson, Audrey Bancod, Valérie Moal, Bernard La Scola

## Abstract

Monocolonal antibodies (mAbs) are currently used for active immunization of COVID-19 in immunocompromised patients. We herein show that in spite there are variations in susceptibility to available mAbs that are authorized for clinical use in France tested on the original B.1.1 virus and 9 variants of concern or of interest, the cocktail casirivimab/imdevimab (REGN-CoV-2) showed a major synergistic effect. However, none of the four mAbs either alone or in combination neutralized the new Omicron variant. Our data strongly warrant a reinforcement of protective measures against infection for immunocompromised patients.

## Text

Monocolonal antibodies (mAbs) are currently used for active immunization of COVID-19 in immunocompromised patients that do not respond to a complete vaccine schedule. As described previously^1^, we tested the neutralizing activity of four mAbs that are authorized for clinical use in France, including bamlanivimab and etesevimab (alone or in combination) and casirivimab and imdevimab (alone or in combination as REGN-CoV-2), against SARS-CoV-2 strains isolated throughout the pandemic. Strains are the French original B.1.1 virus and 9 variants of concern or of interest: B.1.160, Alpha (B.1.1.7), Beta (B.1.351.2), Delta original (AY.71) and of sublineage (AY.4.2), Iota (B.1.526), Epsilon (B.1.429), Mu (B.1.621), and the recent Omicron (B.1.1.529)^2^. Bamlanivimab did not inhibit the Beta and Delta variants as previously reported^3^, but also of Epsilon and Mu variants (Figure 1 and Supplementary appendix). For etesevimab, 50% of neutralization was below 5 µg/mL for Original/B.1.1 virus, Epsilon variant and both delta variants. For the cocktail of bamlanivimab/etesevimab, efficient neutralization was recovered for Alpha, B.1.160 and Iota variants. Casirivimab efficiently neutralized Original/B.1.1 virus, B.1.160, Alpha, Delta, AY4.2, Epsilon and Iota variants. In contrast, we did not observe any neutralization by casirivimab of Beta and Mu variants. Imdevimab neutralized all variants except Omicron but concentrations to obtain 50% of neutralization were higher on average than with casirivimab. Unexpectedly, the cocktail casirivimab/imdevimab showed a major synergistic effect, particularly on Delta, AY4.2 and Epsilon variants because 50% of neutralization was observed at 0.03 µg/mL. We observed 50% of neutralization at 0.2 µg/mL for Original/B.1.1 virus, Alpha and Iota variants, at 0.4 µg/mL for B.1.160, 0.7 µg/mL for Beta. For Mu variant, we observed heterogeneity according to the replicates with 50% of neutralization on average at 2 µg/mL. However, none of the four mAbs either alone or in combination neutralized the new Omicron variant.

**Figure 1:**
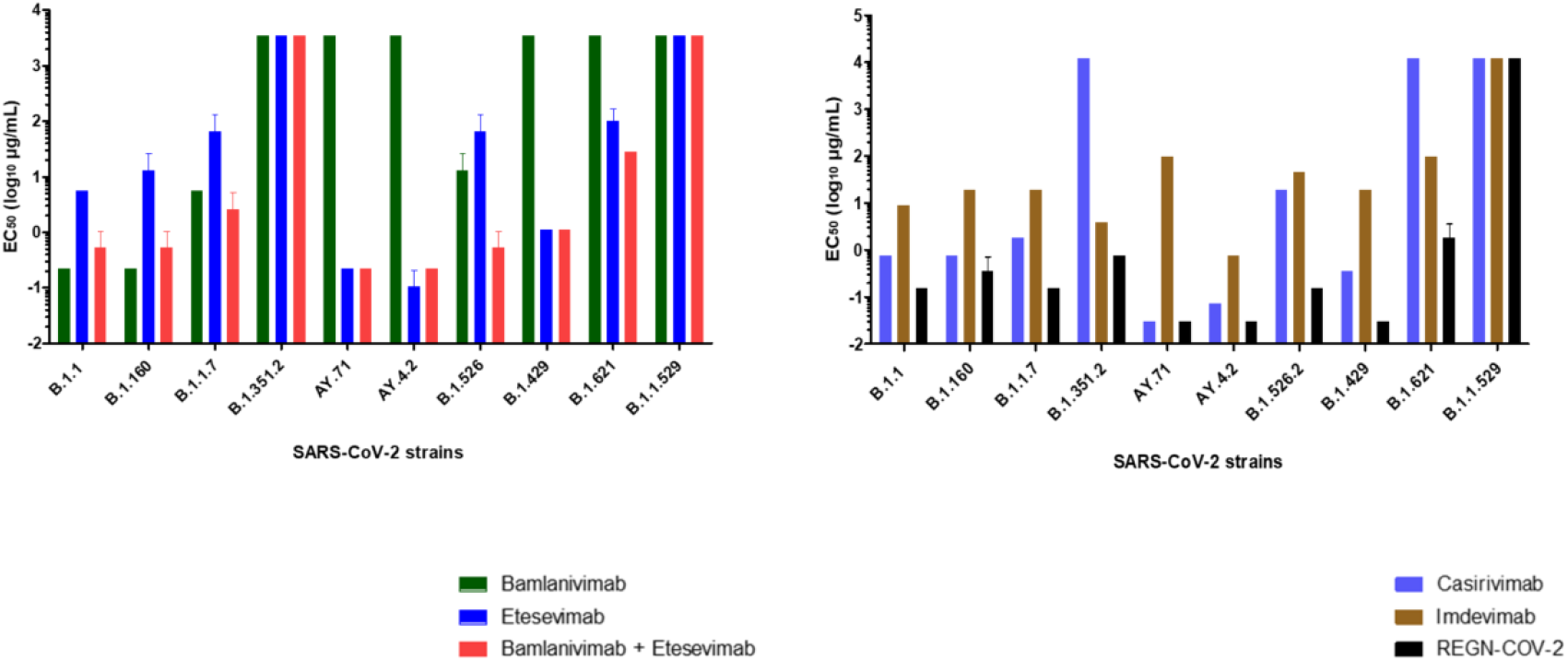
Concentrations required obtaining 50% neutralization (EC_50_ log_10_ µg/mL) for each mAb. (A) bamlanivimab, etesevimab, mixture of bamlanivimab and etesevimab, (B) casirivimab, imdevimab and REGN-CoV-2 on the 10 SARS-CoV-2 strains tested. Each mAb was tested three times (except for Omicron variant 4 times).

These results suggest that although the four tested mAbs can have a lowered effect on recently emerging variants, their combination is highly synergistic *in vitro*, a feature clinically reported recently for Delta variant ^4^. But we observed also that the 4 mAbs currently used alone or in combination in our country showed a complete loss of their neutralizing activity against Omicron variant, a feature recently reported in comparison to the WA1/2020 D614G parental isolate^5^. Of course, definitive conclusions regarding the inefficiency of mAbs against Omicron await the outcomes of clinical studies but our data strongly warrant a reinforcement of protective measures against infection for immunocompromised patients.

## Acknowledgements

the authors are indebted to Pr Philippe Colson for critical review of the manuscript and analysis of genomic data, Audrey Bancod for helping to test microneutralisation of mAbs, Marie-Charlotte Mati and Clio Grimaldier for strains isolation, Gwilherm Penant and Priscilla Jardot for genome sequencing of isolates.

## Author contributions

CB performed microneutralisation tests and co-wrote the first draft of the article, CB performed microneutralisation tests, PC performed genomes analysis co-wrote the first draft of the article, VM co-designed the study, BL conceived and co-designed the study and co-wrote the first draft of the article. All authors contributed to discussion and interpretation of the results, and to the writing of the manuscript. All authors have read and approved the final manuscript.

## Supplementary data

### Materials and methods

#### Cell culture

Vero E6 cells (ATCC-CRL-1586) were cultured without antibiotics in minimal in medium (MEM, Gibco, USA) with 2 mM L-glutamine and 10% foetal bovine serum (FBS) at 37°C in a 5% CO_2_ incubator. Vero E6 cells were then prepared at a concentration of 5×10^5^ cells/mL in ninety-six-well plates for the neutralization tests of SARS-CoV-2 in MEM growth medium with glutamine and 4% FBS (M4 media).

#### SARS-CoV-2 viral strains

The ten SARS-CoV-2 strains used in this study were isolated in cells culture and stored at - 80°C from patients’s nasopharyngal swabs tested SARS-CoV-2 positive in our institute IHU-Méditerranée Infection during the pandemic^1,2^. The supernatant of each strains was then harvested and was genotyped by whole genome next generation (NGS) as previously described^3^ (Supplementary data). For the neutralization tests, we inoculated the viral strains in 96-well Vero E6 cells plate at a concentration of 5×10^5^ cells/mL. Forty eight hours post-viral infection, viral suspension was harvested and quantified by real-time reverse-transcription RT-PCR and TCID50.

#### Monoclonal antibodies dilutions

Bamlavinimab and etesevimab were diluated each in a 1:5 serial dilutions (from 3500 µg/mL to 0.0089 µg/mL). For the combination of the two mAbs, we tested the mixture in the highest concentration for each mAbs alone with 2 times more etesevimab than bamlavinimab. Casirivimab and imdevimab were diluated each in a 1:5 serial dilutions (from 12 000 µg/mL to 0,00614 µg/mL). For the combination of these two mAbs, we tested the mixture in the highest concentration for each mAbs alone in the same proportion.

#### Micro-neutralization assay

Each dilution of mAbs was mixed volume by volume with each viral strains with standardized inoculum at 25 Ct as previously described^4^. The mixture of viral suspension and mAbs was incubated 1h at 37 ° C under 5% CO_2_. Then, 100μl of medium in the 96-well plates was removed and 100μL of the mixture for each dilution was added in quadruplate on the Vero E6 cells. Five days post-viral infection, cytopathic effect was determined with the inverted optical microscope to determine the neutralization titer to obtain 50% of neutralization. Each mAbs and combination of mAbs were tested three times against the 10 SARS-CoV-2 strains, except for omicron variant that was tested four times.

---

**Supplementary figure S1:**
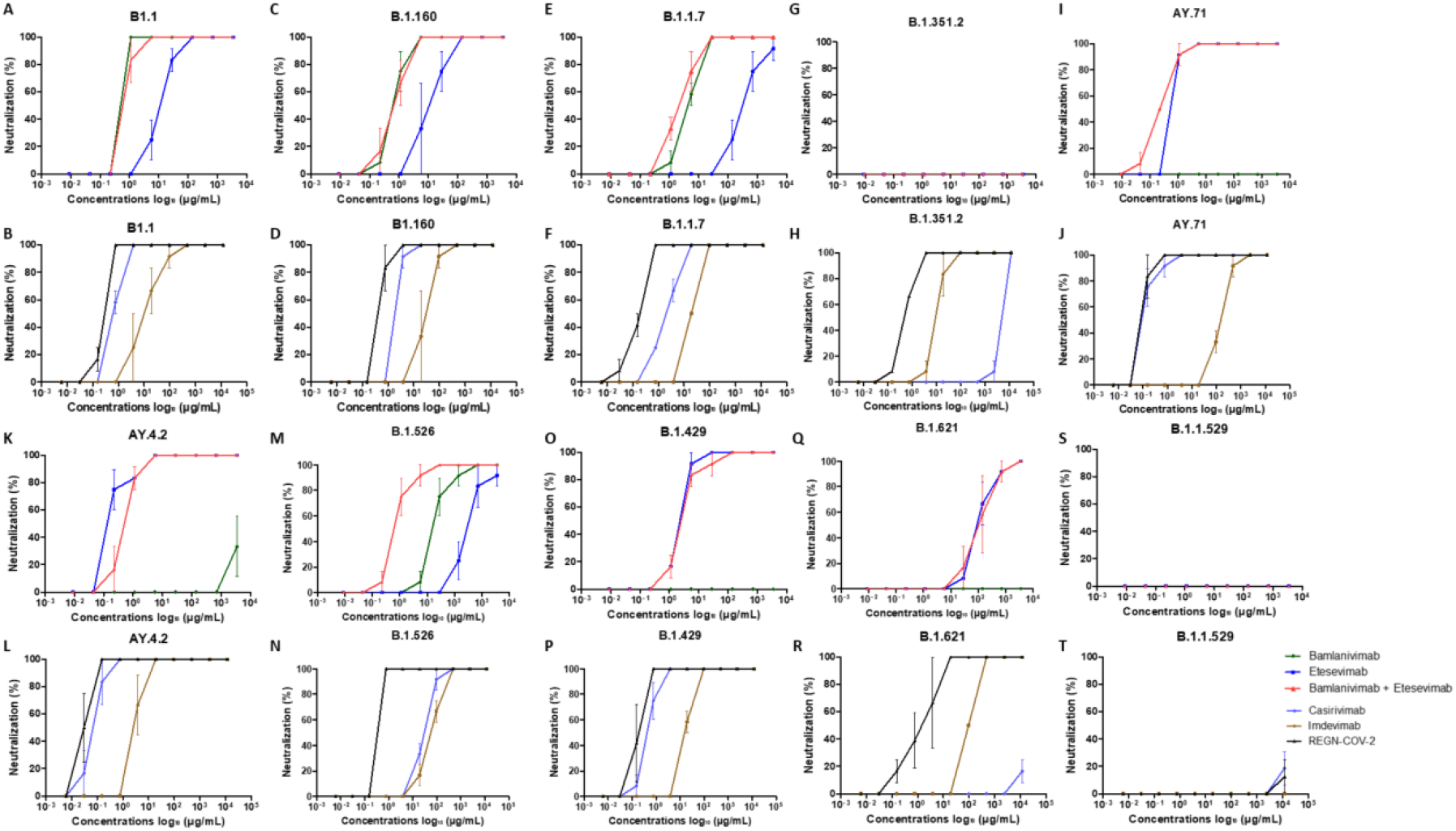
Neutralization curves in Vero E6 cells for each strains tested with each mAb : A, C, E, G, I, K, M, N, O, Q, S : bamlanivimab, etesevimab and mixture of bamlanivimab and etesevimab – B, D, F, H, J, L, N, P, R, T : casirivimab, imdevimab and REGN-CoV-2. Each experiment was done three times, except for Omicron variant four times.

**Supplementary Table S2:**
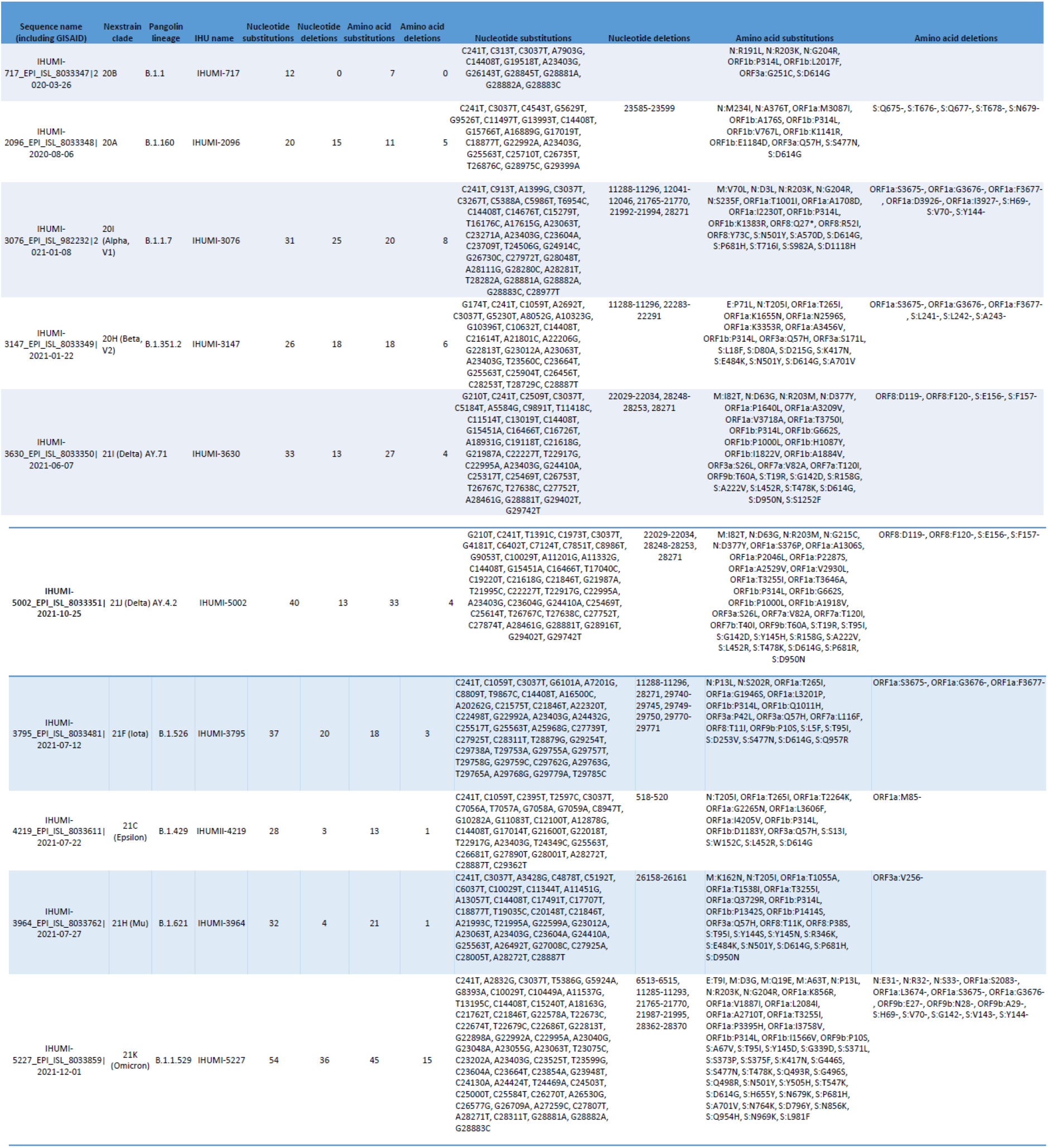
Lineage of SARS-CoV-2 isolates and mutations in the spike protein. In this table are indicated for the ten SARS-CoV-2 strains: genome sequence submitted to GISAID databe (https://www.gisaid.org/), nexstrain clade, Pangolin lineage, IHU name isolate (IHUMI) and the corresponding nucleotide substitutions, nucleotide deletions, amino acid substitutions and amino acid deletion

